# Long-day increase of *HvVRN2* expression marks the deadline to fulfill the vernalization requirement in winter barley

**DOI:** 10.1101/314799

**Authors:** Arantxa Monteagudo, Ernesto Igartua, Ildikó Karsai, M Pilar Gracia, Ana M. Casas

## Abstract

Vernalization and photoperiod cues are integrated in winter barley plants to flower in the right conditions. We hypothesize that there is a timeframe to satisfy the vernalization needs in order to flower in the optimum moment. Growth and expression of different flowering promoters (*HvVRN1*, *HvCO2*, *Ppd-H1*, *HvFT1*, *HvFT3*) and repressors (*HvVRN2*, *HvCO9* and *HvOS2*) were evaluated in two winter barley varieties under: (1) natural increasing photoperiod, without vernalization, and (2) under short day conditions in three insufficient vernalization treatments. Here, we provide evidence of the existence of a day-length threshold, around 12 h 30 min in our latitudes (Zaragoza, Spain, 41°43’N), marked by the rise of *HvVRN2* expression, which defines the moment in which cold requirement must be satisfied to acquire competency to flower. Before that, expression of *HvCO2* was induced and might be promoting *HvFT1* in both inductive and non-inductive conditions. *HvFT3*, to be effectively expressed, must receive induction of cold or plant development, through downregulation of *HvVRN2* and *HvOS2*. We emphasize the contribution of *HvOS2*, together with *HvVRN2*, in the delay of flowering in vernalization-responsive cultivars. Understanding this complex mechanism of flowering might be useful for breeders to define varieties, particularly in a climate change scenario.

## Introduction

Tight coordination of flowering time to environmental conditions is crucial for crop reproductive success and has a major impact on yield (Campoli and von Korff, 2014; Digel *et al.*, 2015). Barley (*Hordeum vulgare* L.) and wheat (*Triticum* spp.) are long-day plants, flowering earlier under increasing day-lengths. Depending on their growth habit, cereals are classified as winter or spring. Winter cereals need a period of exposure to low temperature, a process called vernalization (Laurie *et al.*, 1995; Trevaskis *et al.*, 2003), which must be completed timely so the plant is prepared to take full advantage of the induction of flowering by long days (Trevaskis, 2010). This requirement could make winter barley and wheat more susceptible to climate change, since the probability of accumulating enough cold hours will decrease in warming winters. Winter barley varieties are sown in autumn, benefiting from the warmth of the soils and the humidity from autumn rains, which are essential at the beginning of the cycle. In the Mediterranean region, they have to survive a range of mild to harsh winters, and then flower sufficiently early in the spring to avoid the heat and drought of late spring or early summer.

The accepted gene model for vernalization-responsive varieties establishes that during winter, cold exposure upregulates the floral promoter *HvVRN1*, which is required to downregulate the flowering repressor *HvVRN2*, allowing expression of the flowering inducer *HvFT1* in leaves (Distelfeld *et al*., 2009). *HvVRN2*, *ZCCT-H* gene and member of the *CONSTANS*-like gene family, delays flowering until plants have satisfied its cold needs (Yan *et al.*, 2004). In winter barleys is present in the dominant variant, whose expression is highly dependent on day-length, being induced in long days (Karsai *et al.*, 2005; Trevaskis *et al.*, 2006).

*HvVRN1* encodes an AP1-like MADS-box transcription factor (Danyluk *et al.*, 2003; Trevaskis *et al.*, 2003; Yan *et al.*, 2003). It presents several alleles as a result of deletions or insertions in the first intron, associated with different degrees of vernalization requirement (Hemming *et al.*, 2009). In winter barley, *HvVRN1* is expressed after exposure to low-temperatures (von Zitzewitz *et al.*, 2005; Sasani *et al.*, 2009), although it can be activated by other pathways such as the developmental pathway, with a marked delay compared with induction by vernalization (Trevaskis *et al.*, 2006). Induction of *HvVRN1* is related to changes in the pattern of histone methylation, whose maintenance provides a memory of cold exposure in winter barley plants (Oliver *et al.*, 2009).

Under short day (SD) conditions, *HvFT3*, a FT-like member of the PEBP family, and candidate gene for *Ppd-H2* (Faure *et al.*, 2007; Kikuchi et al., 2009), has been described as promoter of flowering (Laurie *et al.*, 1995). Two allelic variants for this gene are known: a dominant one, with a functional copy of the gene, and a recessive allele, with most of the gene missing and nonfunctional (Kikuchi *et al.*, 2009). Its presence caused differences in heading date in SD (Boyd *et al.*, 2003; Faure *et al.*, 2007). A QTL for heading date co-locating with this gene was identified in autumn sowings (Cuesta-Marcos *et al.*, 2008; Borràs-Gelonch *et al.*, 2012), showing a remarkable importance for adaptation under Mediterranean conditions (Casao *et al.*, 2011*b*), although it may have a negative impact for low temperature tolerance in facultative genotypes (Cuesta-Marcos *et al.*, 2015, Rizza *et al*. 2016).

The sensitivity to long days (LD) is determined by *Ppd-H1* (Laurie *et al.*, 1995), candidate gene for *HvPRR7*, pseudo-response regulator 7 (Turner *et al.*, 2005). The dominant allele accelerates flowering mediating the induction of *HvFT1* through the activity of *HvCO1/HvCO2* (Turner *et al.*, 2005). Recently, Mulki and von Korff (2016) have revealed a possible role of *Ppd-H1* as repressor of flowering, mediating the induction of *HvVRN2* before the vernalization

Another known repressor in the vernalization pathway is the monocot homolog of the Arabidopsis *FLOWERING LOCUS C*, *ODDSOC2* (in barley *HvOS2*). It is a MADS-box repressor of flowering, also downregulated by vernalization (Greenup *et al.* 2010; Ruelens *et al.* 2013), which is affected by photoperiod and induced by high temperatures (Hemming *et al.*, 2012).

VRN1 directly binds to the promoter regions of the repressor genes *HvVRN2* and *HvOS2*, downregulating their expression, and also to the *HvFT1* promoter, enhancing its expression (Deng *et al.*, 2015). These results explain why vernalization is a pre-requisite to promote flowering under long-day (LD) in temperate cereals. However, it has been suggested that other additional genes may be acting as regulators of *VRN2* when exposed to cold (Chen and Dubcovsky, 2012; Sharma *et al.*, 2017).

Two closely related *CO* genes, *CO1* and *CO2*, are present in the temperate cereals. Both are LD-flowering promoters modulated by circadian clock and day-length (Griffiths *et al.*, 2003; Nemoto *et al.*, 2003). CO2 competes with VRN2 for interactions with a common protein, NF-Y, in a mechanism to integrate environmental cues through regulation of *HvFT1* (Li *et al.*, 2011). Overexpression of *HvCO2*, induced the expression of *HvFT1* in spring barley but it caused up-regulation of the repressor *HvVRN2*, in winter barley, resulting in reduced expression of *HvFT1* and delayed flowering (Mulki and von Korff, 2016). Other member of the *CONSTANS-*like family, *HvCO9* (*HvCMF11*,Cockram *et al*. (2012)) is a negative regulator of flowering, paralogue of *HvVRN2* (Higgins *et al.*, 2010). It is expressed under non-inductive SD conditions, correlating with *HvFT1* and *HvFT2*, but no relationship was found between *HvCO9* and *HvFT3* (Kikuchi *et al.*, 2012).

Crop modelling studies have determined that barley ideotypes, for future Boreal and Mediterranean climatic zones in Europe, should have appropriate vernalization and photoperiod responses finely tuned to the needs of each specific region (Tao *et al.*, 2017). One future avenue for plant breeding will be to use elite germplasm coming from regions that have experienced the foreseen conditions (Atlin *et al.*, 2017). For example, transferring cultivars adapted to Mediterranean conditions, which possess a strategy based on scape to drought, to more northern latitudes. To achieve that goal, the responses to photoperiod should be modified accordingly to avoid yield penalties (Dawson *et al.*, 2015). For this reason, comprehensive studies on the effect of photoperiod on major flowering genes are called for.

This study focuses on the repression of flowering, under non-inductive conditions, in winter barley. Previous studies have demonstrated that *HvVRN2* expression needs induction by long days (Trevaskis *et al.*, 2006). But, how long? As studies have been performed under fixed photoperiods in growth chambers, the actual day-length threshold to induce *HvVRN2* is unknown. There is indication that this gene has no effect below 12 h (Karsai *et al*. 2006). This question is relevant from the agronomic point of view. We hypothesize that there is a vernalization window for satisfying the cold requirement, in order to make the plant competent to flower at the right time and achieve a good yield. In these experiments, the goal was to determine the day-length threshold leading to induction of the repressor *HvVRN2*, assuming that this is the end of that window. Additionally, we want to further characterize the role of *HvFT3* in the promotion to flowering in winter barley. Here, we investigate the effects of photoperiod on the transcript levels of selected genes in winter barley, by examining photoperiod responses in the medium-long term (21 - 90 days). The results provided may help to understand the complex mechanism of flowering in suboptimal conditions, and facilitate breeding for present and future climate conditions in Europe and elsewhere.

## Materials and methods

### Plant materials

Two French winter barley varieties, ‘Hispanic’ (two-rowed, ‘Mosar’ x (‘Flika’ x ‘Lada’)) and ‘Barberousse’ (six-rowed, (‘Hauter’ x (‘Hatif de Grignon’ x ‘Ares’)) x ‘Ager’) were selected. They have the same allelic combination in *HvVRN1* (winter allele, same first intron length), *HvVRN2* (all *ZCCT-H* genes present), and *Ppd-H1* (dominant, long photoperiod responsive), but differ in *HvFT1* and *HvFT3* (*Ppd-H2*, present in ‘Hispanic’, defective in ‘Barberousse’) (Loscos *et al.*, 2014).

### Plant growth, phenotyping and sampling

#### Experiment 1 – Sowings under increasing natural photoperiod

For each variety, we used two 1L-pots at each sowing time (standard substrate made of peat, fine sand and perlite, from a mix with 46 kg, 150 kg and 1L, respectively). Pots were sown with 7 seeds once a week, sequentially, from Feb 11^th^ until April 8^th^ 2015, in a glasshouse in Zaragoza (41°43’N, 00°49’W) under natural photoperiod (Fig. 1) and controlled temperature (22±1°C day / 18±1°C night). Unless specified, plants were not vernalized (NV). Spatial homogeneity in irradiance was obtained rotating the plants each week. As vernalized control, three pots of each variety were sown on Feb 11^th^. They were grown during 7 days (until germination) under glasshouse conditions, and then were vernalized (VER) under short photoperiod (8 h light) and 6±2°C for 49 days. After the cold treatment, plants were transferred to the same glasshouse on April 8^th^, when natural photoperiod was 13 h. Duration of daylight at sowing and sampling dates was gathered from http://www.timeanddate.com/sun, taking sunrise and sunset as the times when the upper edge of the Sun’s disc touches the horizon.

**Figure 1.**
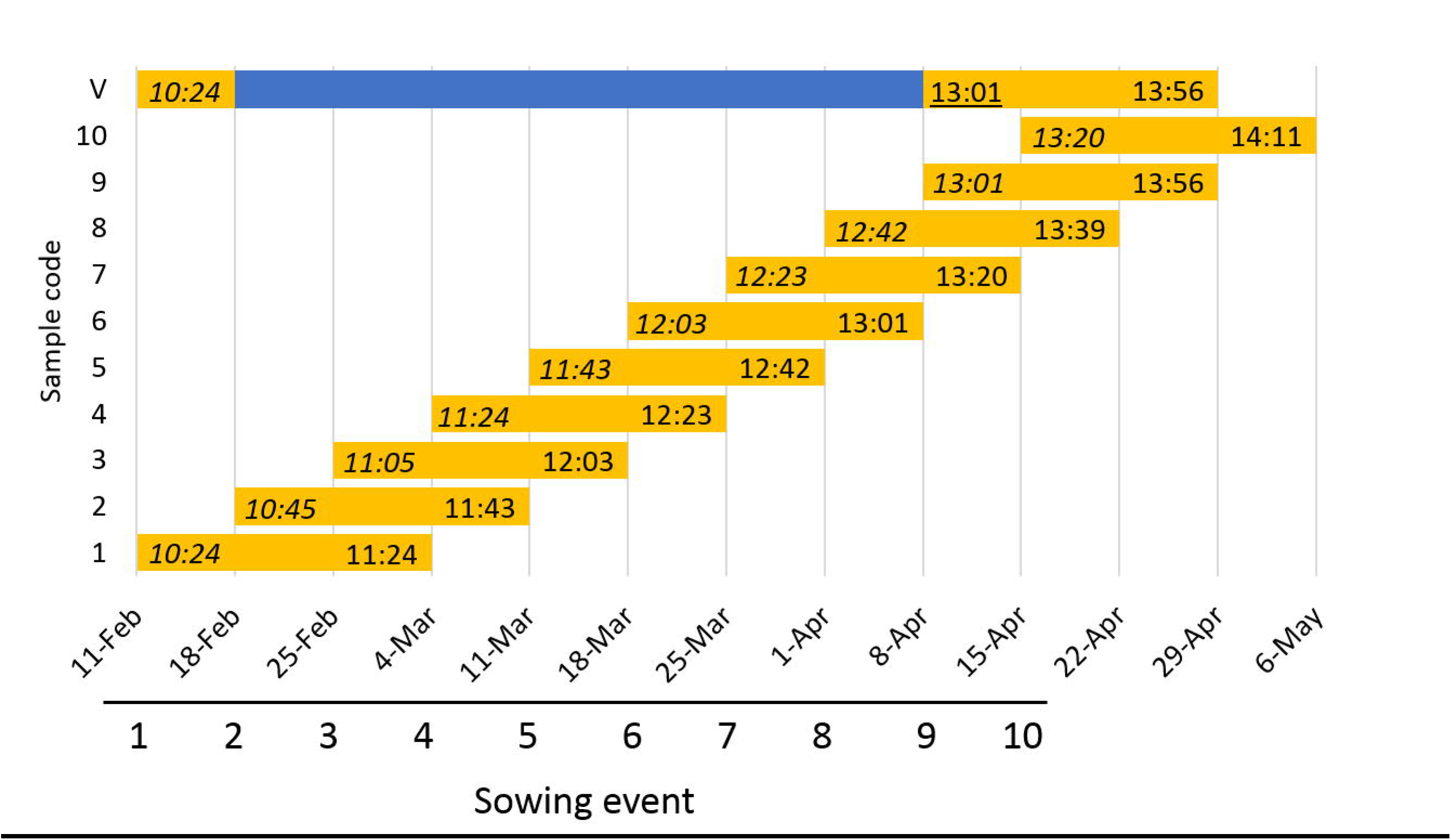
Experiment planning. Each sowing and its sampling are represented. Yellow bars show the time that plants were kept under non-vernalized conditions. Blue bar shows the time spent in the vernalization chamber. X-axis represent dates of start of experiment and sampling date (three weeks after sowing). The second numbers inside the yellow bars are the day-length at sampling date (HH:MM). The first numbers in italics represent day-length at sowing day, and underlined numbers are day-length in the shift day (vernalized plants were transferred to glasshouse). Sunrise and sunset are the times when the upper edge of the Sun’s disc touches the horizon.

For gene expression, the last expanded leaf of three 21-day-old plants (3-leaf stage) was sampled 8 h after dawn, frozen in liquid nitrogen, homogenized (Mixer Mill model MM301, Retsch) and conserved at −80°C until RNA isolation. In all the experiments, sampling time was chosen to capture high expression moments of almost of the genes involved, taking into account their circadian rhythms.

On a fixed date (19^th^ May, day-length 15 h, 97 days after the first sowing), we took a cross-sectional sample across sowing events. The last expanded leaf of each weekly-sown plant was sampled 12 h after dawn for RNA isolation. Then, dissection of the plants (all stems of each plant) was made in order to determine the development of the apex (with naked eye, reproductive apex was equivalent to more than 3 mm).

#### Experiment 2 – Growth chamber, 12 h light

Seventy-two seeds of each variety were sown in 12-well trays (650 cc) and allowed to germinate during 7 days in a growth chamber at 12 h light, 20°C/12 h dark, 16°C, 65% HR and light intensity of 300 μmol m^-2^s^-1^ PAR. Then, the trays were divided in three groups that received the following treatments: (A) NV, (B) 14-days VER and (C) 28-days VER. Group A stayed at the growth chamber while B and C were transferred to a vernalization chamber, 8 h light/16 h night and constant temperature (6 ± 2°C). Groups B and C were returned to the growth chamber after 14 and 28 days of cold treatment, respectively. After forty days at the growth chamber, 3 plants of each variety and treatment were transferred to a 1L pots to let them grow until flowering.

For gene expression the last expanded leaf of four plants was sampled 14, 28, 35 or 49 days after germination (A) or after the end of the VER treatment (B and C), 10 hours into the light period.

Number of leaves, tillers, development at Zadoks scale (Zadoks *et al.*, 1974) were recorded along the experiment every 3-5 days. LSD multiple comparisons were obtained for each trait. Also apex dissections were carried out at selected time points to establish the Waddington developmental stage (Waddington and Cartwright, 1983). The experiment ended 136 days after sowing.

### Vernalization response of ‘Hispanic’ and ‘Barberousse’

In the course of earlier experiments, carried out in the Phytotron of Martonvásár (Hungary), both varieties were exposed to different VER treatments (0, 15, 30 or 45 days, 4°C, 8 h light), and then transferred to a growth chamber 16 h day-length, 18°C and light intensity of 340 μmol m^-2^ s^-^1. Flowering date was recorded at each treatment (Fig. S1).

### Gene expression analysis

Three individual plants were sampled at each time point per genotype. RNA extraction was carried out using Nucleospin RNA Plant Kit (Macherey-Nagel, Germany) following manufacturer instructions. Total RNA (1μg) was employed for cDNA synthesis using SuperScript III Reverse Transcriptase (Invitrogen) and oligo (dT)_20_ primer (Invitrogen). Real-time PCR quantification (ABI 7500, Applied Biosystems) was performed for samples from each time point from NV plants and for VER plants as control treatment.. Three biological repeats and two technical repeats were performed per sample and pair of primers (*HvVRN1*, *HvVRN2*, *Ppd-H1*, *HvCO2*, *HvCO9*, *HvOS2, HvFT1*, and *HvFT3*). Primer sequences and conditions are specified in Table S1. Expression levels were normalized to *Actin* expression, taking into account primer efficiencies.

### Statistical analysis

Statistical analysis was carried out in R software (R Core Team, 2017). Multiple comparisons were obtained by Fisher’s protected Least Significant Differences (LSD) with the R package ‘agricolae’ (de Mendiburu, 2016). For gene expression results, the mean of two technical replications of ∆Ct (Ct actin – Ct target) was used as unit. Analyses of variance for each gene and experiment were performed considering all factors as fixed. Pearson correlations were carried out with ‘cor’ function.

## Results

### Gene expression under increasing photoperiod conditions (experiment 1)

Natural day-length at sampling increased from ~11 h 30 min at the first sowing to ~14 h at the 9^th^ sowing event, and also for the VER control (Fig. 1).

Leaves were sampled 21 days after each sowing or 21 days after the end of the VER treatment. Expression levels of *HvCO2*, *HvCO9*, *HvFT1*, *HvFT3*, *HvOS2*, *HvVRN1*, *HvVRN2* and *Ppd-H1*, were analysed by qRT-PCR. *HvVRN1* expression was detected only in VER plants (Fig. 2), in concordance with the cold treatment received. *HvVRN2* expression was detected at all time points, although expression was lower in plants from the first four sowings, grown under shorter photoperiods (Fig. 2). Concurrent higher levels of *HvCO2* were detected in those same plants (Fig. 2). When the day-length reached 12 h 30 min (sowing events 5-9), corresponding to 28^th^ March in our latitudes, the expression of *HvCO2* decreased, *Ppd-H1* increased in ‘Hispanic’ and the levels of *HvVRN2* increased in both genotypes.

**Figure 2.**
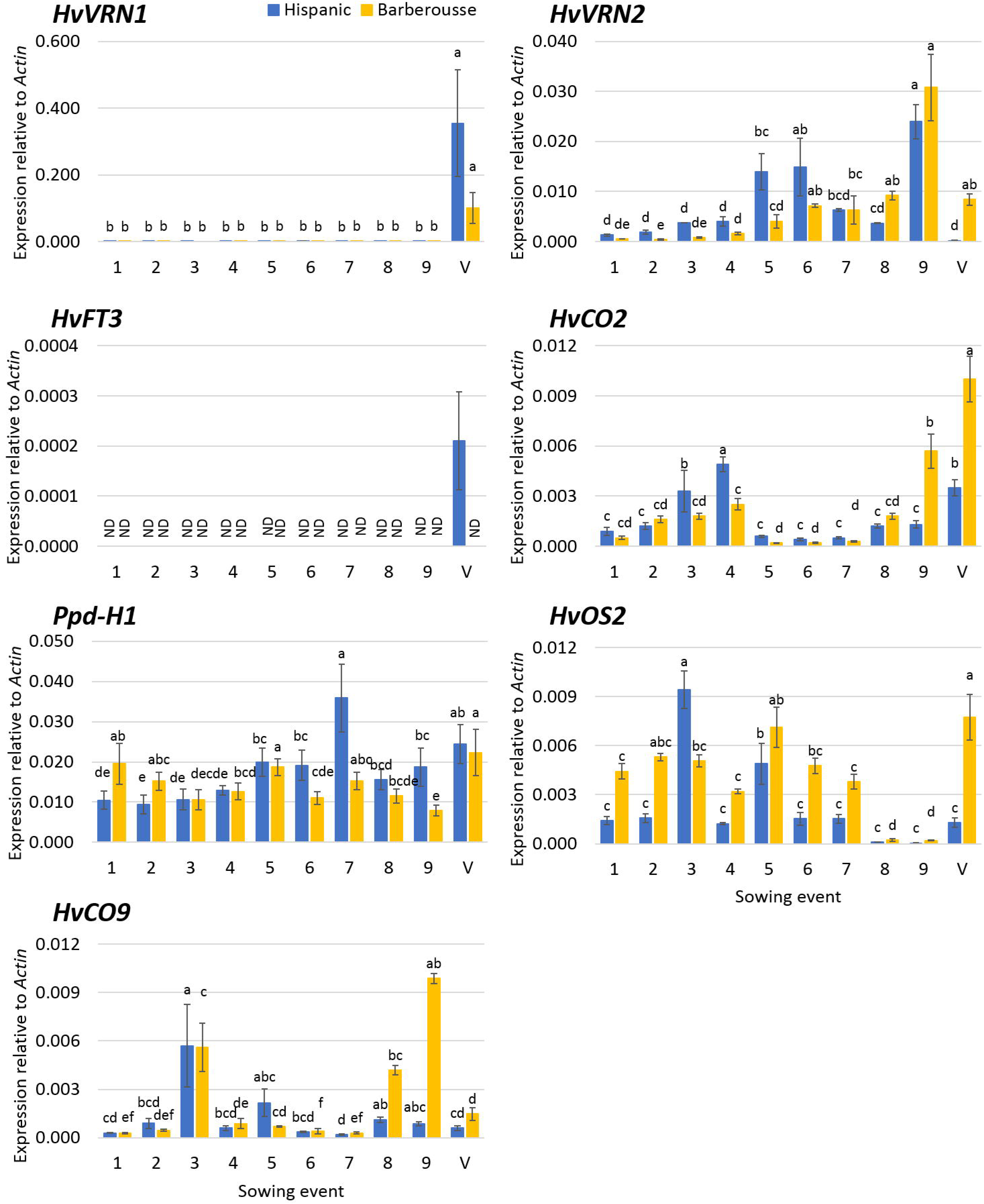
Gene expression three weeks after sowing. X-axis represent the successive sowings, from 11th February until 8th April. Unvernalized plants (sowings 1 to 9) and vernalized control (V) of ‘Hispanic’ (blue) and ‘Barberousse’ (yellow) are plotted. Mean of 3 biological replicates. Error bars are SEM. ND, Not detected. *HvFT1* expression is not reported as it was null for all non-vernalized samples. For each variety, bars with the same letter are not significantly different at P<0.05 (LSD test).

Therefore, upregulation of *HvVRN2* in NV plants occurred under day-lengths longer than 12 h 30 min in 21 day-old plants, and was also detected in 14 day-old plants (Fig. S2).

Without vernalization, neither genotype showed expression of *HvFT3* (Fig. 2). This was expected for ‘Barberousse’, as it has the null allele, but we could not anticipate this result for ‘Hispanic’. In this genotype, the expression levels were below the detection limit, except for VER plants.

In general, ‘Barberousse’ presented higher *HvOS2* expression levels than ‘Hispanic’, except for the last samplings, when *HvOS2* expression was barely detectable in both genotypes. Expression of *HvCO9* in ‘Hispanic’ was low and variable. Higher *HvCO9* expression was observed in the last time points of ‘Barberousse’ (Fig. 2).

Differences between genotypes were also detected in VER plants. High expression of *HvVRN1* and *HvFT3*, and barely any expression of *HvVRN2*, was seen in ‘Hispanic’. On the other hand, although *HvVRN1* was detected in VER plants from ‘Barberousse’, high expression of *HvOS2* and *HvVRN2* was apparent (Fig. 2), suggesting a delay in development, which was even more evident when assessing apex growth (Fig. 3).

**Figure 3.**
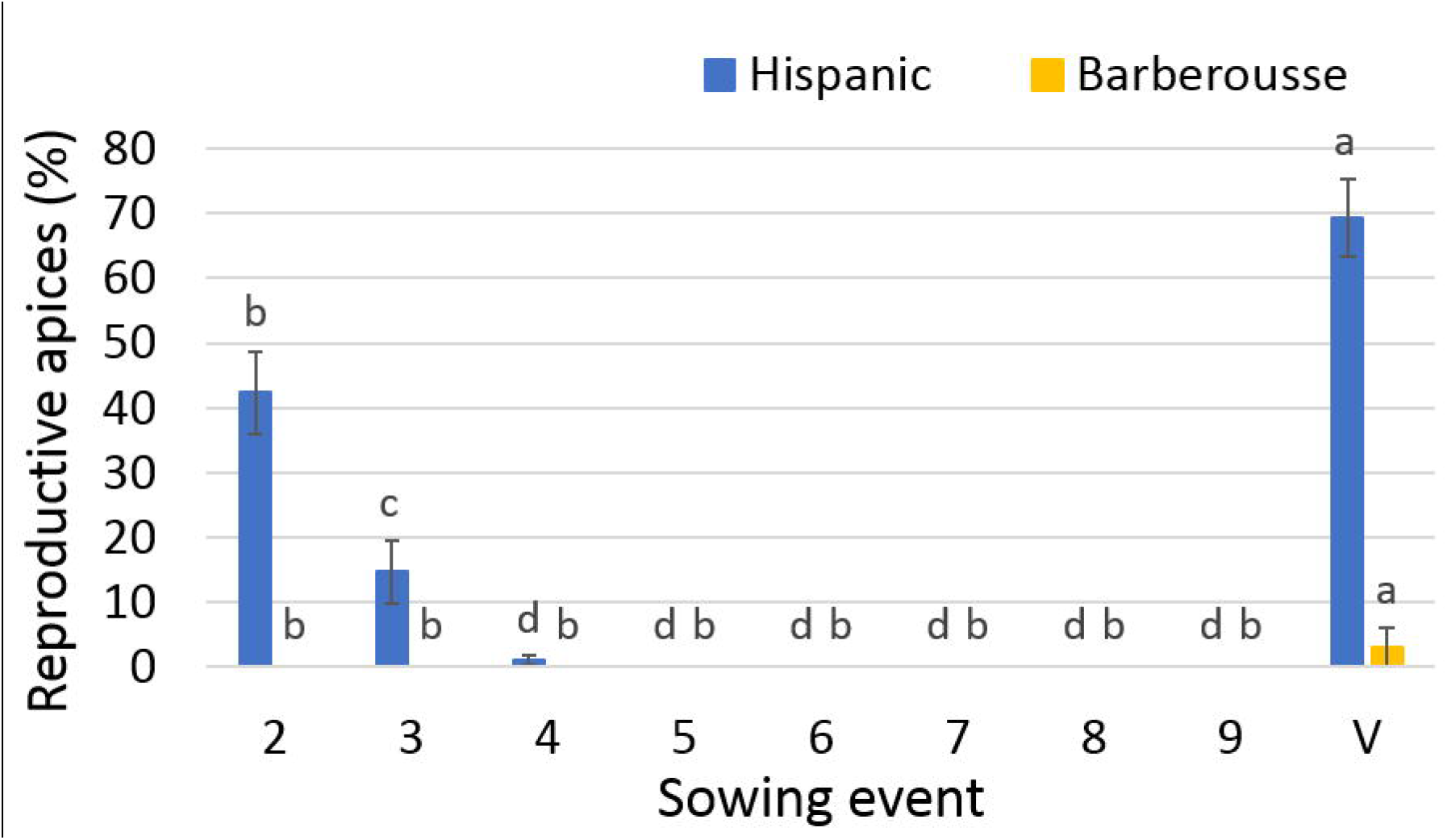
Percentage of reproductive apices regarding the total (vegetative and reproductive) after 100 days of the experiment. Mean of 10-12 plants. Error bars are SD. For each variety, bars with the same letter are not significantly different at P<0.05 (LSD test).

### Reproductive stage and gene expression from medium to long term development at 15 h day-length (experiment 1)

Ninety-seven days since the beginning of the experiment, on **May 19^th^**, with 15 h light, the number of apices at reproductive stage per plant was recorded (Fig. 3). Among NV plants, only the second sowing event of ‘Hispanic’ reached the stage Z49 (first awns visible) at the end of the experiment (83 days after sowing; no data available for the first sowing, whose plants were dissected earlier and showed reproductive apices after 72 days). VER ‘Hispanic’ and ‘Barberousse’ also showed apices at reproductive stage, ‘Barberousse’ more delayed than ‘Hispanic’.

Expression levels on this same date were also analysed (Fig. 4). Under NV conditions, flowering promoters (*HvVRN1*, *HvFT1* and *HvFT3*) were induced only in ‘Hispanic’ at the first point available (sowing event 2), and were absent in ‘Barberousse’. Accordingly, *HvVRN2*, *HvCO9* and *HvOS2* were repressed in ‘Hispanic’, and induced in ‘Barberousse’. *Ppd-H1* was expressed at higher levels in ‘Hispanic’ and only *HvCO2* was equally expressed in both varieties at this point.

**Figure 4.**
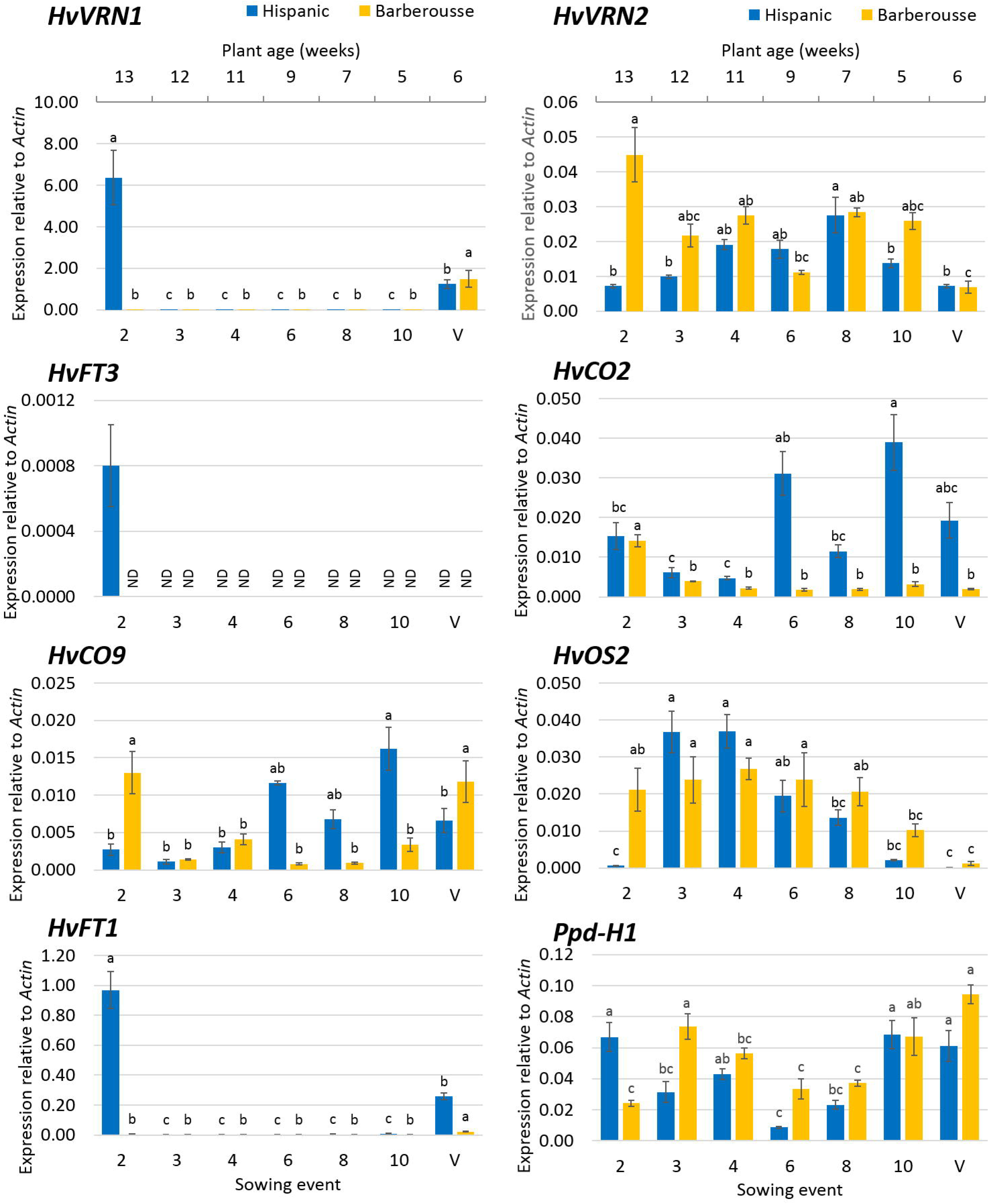
Cross-sectional gene expression under 15 h of natural daylight of the sequential sowings under natural photoperiod experiment. X- upper axis represent the weeks after sowing of unvernalized plants. Control plants (V) were maintained under natural photoperiod for 6 weeks after 49 days of vernalization. Enlarged view of the *HvFT1* expression is shown. Mean of 3 biological replicates. Error bars are SEM. For each variety, bars with the same letter are not significantly different at P<0.05 (LSD test).

For the rest of NV plants (Fig. 4, sowing events 3 to 10), no expression of *HvVRN1*, *HvFT1* or *HvFT3* and high expression of *HvVRN2* and *HvOS2* was detected. Differences between varieties were found in *HvCO2* and *HvCO9* expression, being low in ‘Barberousse’ from sowing events 3 to 10, while levels increased from sowing event 6 in ‘Hispanic’, revealing a common effect of development and variety for these two *CONSTANS*-like genes. Contrasting with this, VER plants did not show differences in transcript levels, except for *HvCO2* and *HvFT1*, which were more expressed in ‘Hispanic’, and less in ‘Barberousse’.

### Responses to 12h photoperiod after increasing vernalization treatments (experiment 2)

Experiment 1 made evident that gene expression was dependent of the plant’s developmental stage (Fig. S2). Two weeks after sowing was not enough to observe differences, but 3 weeks was (Fig. 2). Therefore, for some genes, induction was dependent on plant age. A second experiment was conceived, to assess the relevance of other factors on gene expression, namely day-length, plant age and degree of vernalization. Thus, we set the day-length at 12h, representative of day-length around the start of stem elongation in natural conditions in our region, and short enough not to elicit LD responses. This was combined with insufficient vernalization.

Flowering time was advanced in an inversely proportional manner to the duration of the VER treatment (Fig. 5). Under 12 h of light, and NV, ‘Hispanic’ reached awn tipping (DEV49) after 124 days, whereas ‘Barberousse’ did not reach that stage during the entire experimental period (136 days). Two or four weeks of VER decreased markedly the time to DEV49 for both genotypes. Plants from both VER treatments reached this stage before the NV plants did. Most of this shortening occurred in the period until first node appearance (DEV31), although some additional acceleration was observed between DEV31 and DEV49. Under the conditions of experiment 2, ‘Hispanic’ had clearly higher total and reproductive tillers than ‘Barberousse’ (Fig. 5), increasing with the length of the VER treatment.

**Figure 5.**
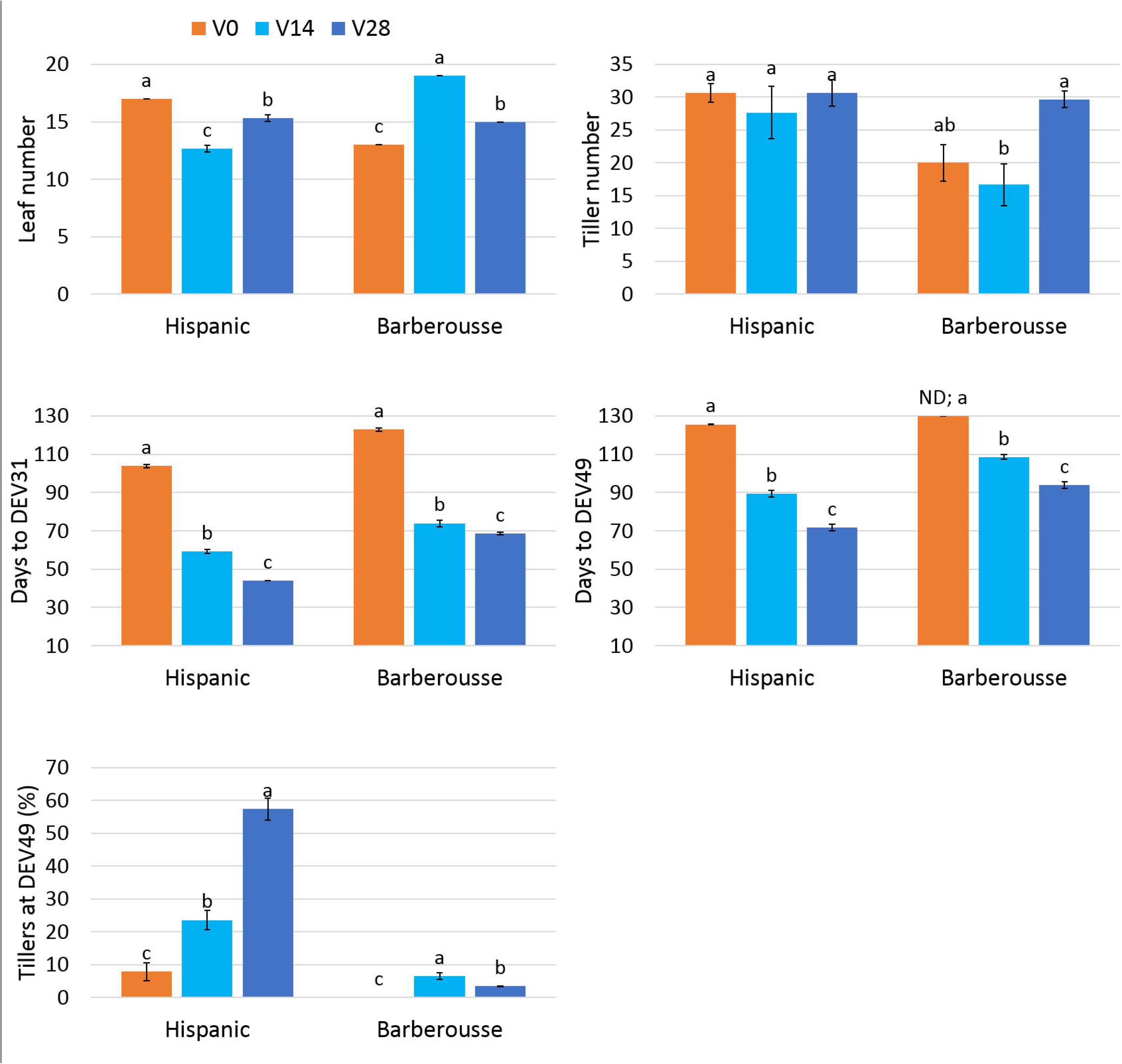
Phenotype of plants growing under controlled conditions (12 h daylight) after different vernalization treatments. Mean of 3 plants. Error bars are SD. ND, no detected (discarded after 130 days without flowering). For each variety, bars with the same letter are not significantly different at P<0.05 (LSD test).

### Gene expression under photoperiod of 12 h affected by vernalization and plant age

Expression analysis showed higher *HvVRN1* induction with the VER duration in both varieties (Fig. 6). Concurrently to the larger expression of *HvVRN1, HvVRN2* was repressed, as expected. Expression of *HvCO9* and *HvOS2* was also reduced with increasing VER. These three repressors showed higher levels in ‘Barberousse’ than in ‘Hispanic’ (Fig. 6), which were correlated with the delayed flowering of ‘Barberousse’ (Fig. S3a). Transcript levels of *Ppd-H1* were similar between treatments. Only ‘Hispanic’ V28 and ‘Barberousse’ V0 showed differences between sample points. Expression of *HvCO2* was clearly related to that of *HvFT1* in both genotypes, showing ‘Barberousse’ earlier induction but lower expression levels (Fig. 6). Such decrease in ‘Barberousse’ is simultaneous with an increased expression of *HvVRN2*, *HvOS2* or *HvCO9*. *HvFT3* transcript levels were present in ‘Hispanic’, only after plants where 28-days VER, and concurrent with a total absence of *HvVRN2*. The decreased expression of *HvCO2* at the last sampling point was inversely related with the longer VER treatment and the early flowering (Fig. S3b).

**Figure 6.**
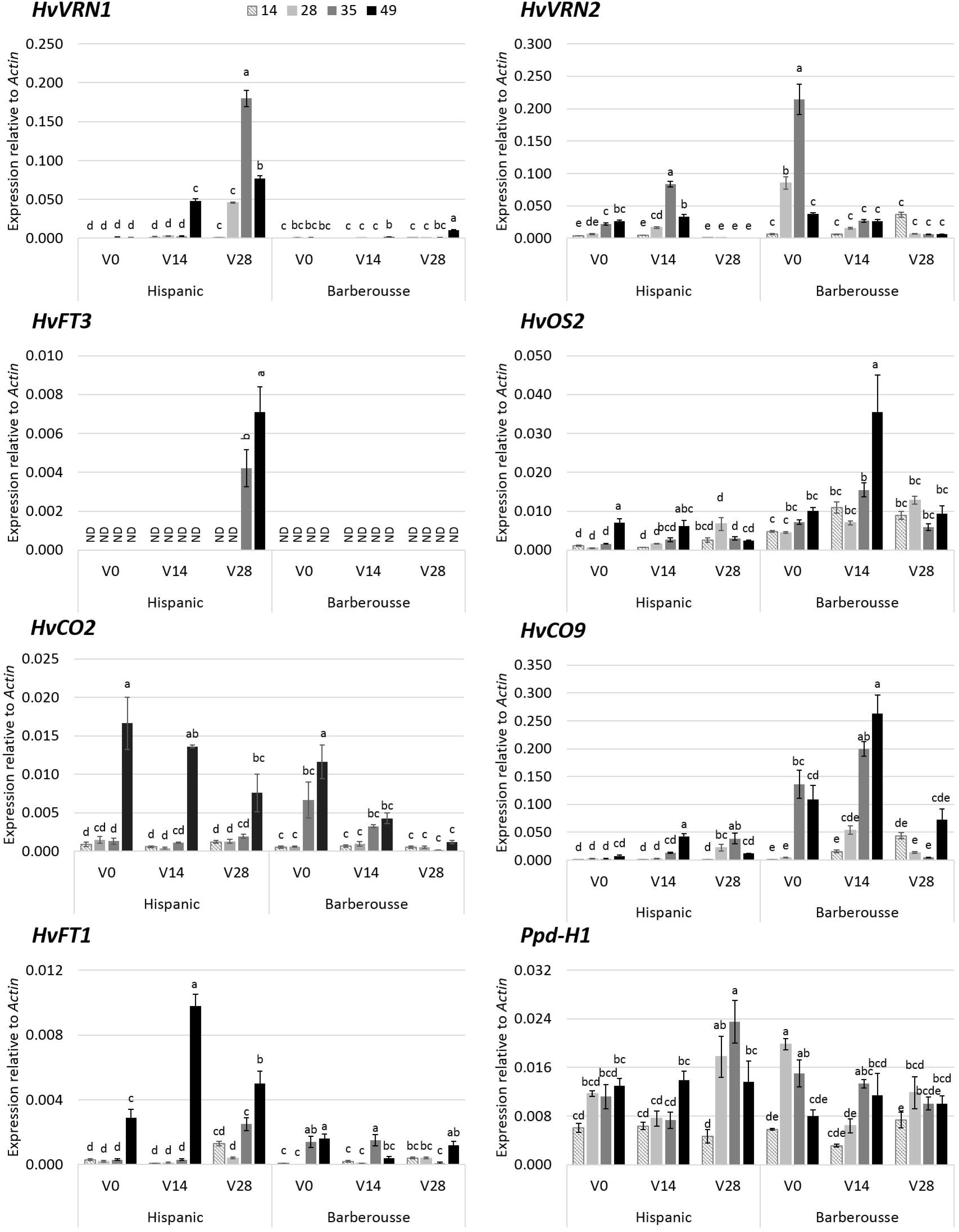
Gene expression under 12 h daylight in growth chamber. X-axis represent days of vernalization chamber. Increasing grey scale is the days after the end of the vernalization treatment when leaves were sampled (14, 28, 35 or 49 days). Mean of 3 biological replicates. Error bars are SEM. For each variety, bars with the same letter are not significantly different at P<0.05 (LSD test).

The increased expression levels of the flowering promoters and the decreased levels of the flowering repressors (Fig. 6) agree with the transition from vegetative to reproductive stage (from W2 to W3; Fig. 7), which was observed only in ‘Hispanic’ VER 28 days. In contrast, ‘Barberousse’ apices only reached this stage unless vernalized and later in time (Fig. S4).

**Figure 7.**
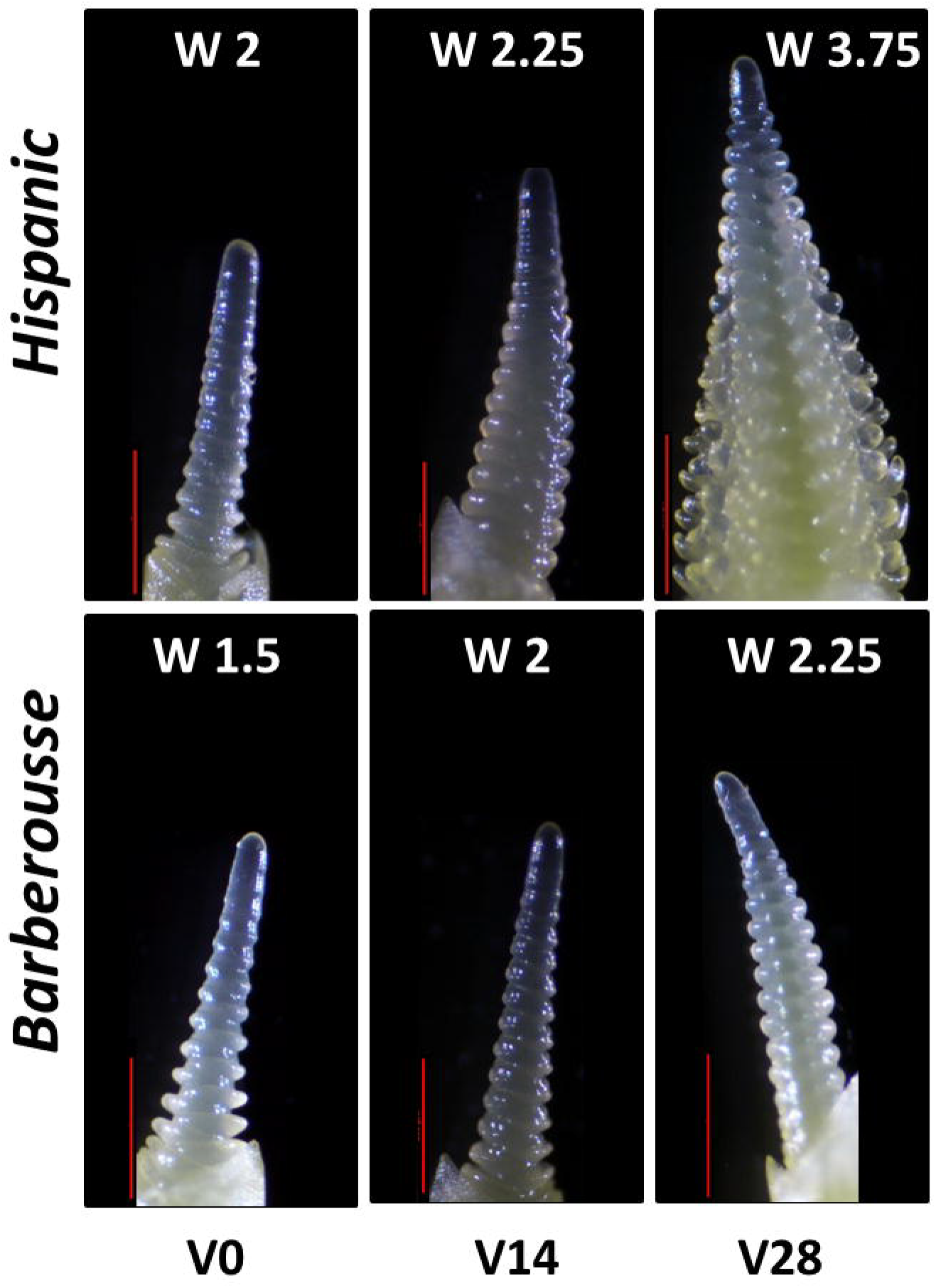
Apex dissection of plants grown under 12h light, 4 weeks after each vernalization treatment. Red bar is 500 μm.

## Discussion

Our results shed further light on the functioning and integration of the vernalization and photoperiod pathways in barley. Although the experiments were performed under controlled conditions, these were established bearing in mind their representativeness of natural conditions. The combined results of the two experiments led us to put forward several new hypotheses to explain the dynamics of vernalization in winter barleys. This and other studies, show that the vernalization process involves more genes than previously thought. The complexity found is challenging but, on the other hand, widens the catalogue of genes amenable for breeding.

### Expression of *HvVRN2* is upregulated beyond 12 h 30 min natural daylight in absence of vernalization

Under typical autumn sowings, winter barley is capable of responding to long photoperiods only after fulfilling a variety-specific low temperature requirement. The model described by Distelfeld *et al.* (2009) and Trevaskis (2010) suggests that, flowering of autumn-sown cereals is delayed before winter because neither day-length nor vernalization response pathways are active. Under SD and low temperature conditions, *HvVRN1* is gradually induced and represses *HvVRN2* to promote flowering. There is also evidence that expression of *HvVRN2* is upregulated in LD (16 h light) and downregulated in SD (8 h light) to almost complete repression (Trevaskis *et al.*, 2006) but, the environmental threshold that induces expression of *HvVRN2* was still undefined. Karsai *et al.* (2006) found a QTL co-locating with *HvVRN2* in vernalized plants when day-length was over 12 hours. Therefore, the limit is similar, irrespective to vernalization.

We found expression of *HvVRN2*, even if at low levels, in NV plants under natural SD (sowings 1-4 in experiment 1). Gene expression remained low until a sudden surge around sowing event 5 (Fig. 2), coincident with an increase of natural daylight above 12 h 30 min (~28^th^ March). We propose that this rise marks the day-length threshold, defining the moment in which cold needs must be satisfied, to acquire competency to flower timely, or else high *HvVRN2* levels will delay flowering beyond agronomically acceptable. This hypothesis should be put to test with specific field experiments.

### Expression of *HvVRN2* does not cause genotypic differences in earliness among two winter genotypes

The comparison of the two winter cultivars clearly revealed a faster early development of ‘Hispanic’, although they are similarly responsive to VER (Fig. S1). Differences in *HvVRN2* expression cannot be the cause of earliness differences, as the early genotype has larger transcript amounts of the repressor. This indicates that there are additional earliness factors differentiating ‘Hispanic’ and ‘Barberousse’, affecting apex development. Such factors would be needed to counteract the repressing effect caused by upregulated *HvVRN2* expression levels. We have explored the possibility that this factor is *HvFT3*, candidate for *Ppd-H2*, the gene affecting short photoperiod sensitivity (Laurie *et al.*, 1995).

### *HvFT3* expression needs induction by cold and plant development, through downregulation of flowering repressors, under non-inductive conditions

The two varieties differ (among others) in the presence/absence of *HvFT3*, which could be the key factor that differentiates them. This gene bears particular agronomic relevance for Mediterranean environments, as it stands at the peak of flowering time QTL and grain yield QTLxEnvironment peaks in several populations (Cuesta-Marcos *et al.*, 2008, 2009; Karsai *et al.*, 2008; Francia *et al.*, 2011; Tondelli *et al.*, 2014). A supporting role for promotion to flowering in winter cultivars, receiving less than enough vernalization under field conditions, was proposed for *HvFT3* (Casao *et al*. 2011*b*). Its expression is usually reported in SD, although it is also found in LD conditions (Kikuchi *et al.*, 2009; Casao *et al.*, 2011*a*). In our experiments, under natural photoperiod, *HvFT3* transcripts were only detected: (a) after full or partial vernalization, in early-medium development (Figures 2 and 6), and (b) in absence of vernalization, in rather late developmental stages, and only in plants sown under shortest day-lengths (Figure 4). We expected expression of *HvFT3* at least in the earliest sowings, experiencing the shortest photoperiods. Instead, it was effectively repressed, either by the low but always presence of *HvVRN2*, or by other repressors. Under constant photoperiod of 12h, *HvFT3* was detected in ‘Hispanic’ only after four weeks VER (2 weeks were insufficient) and 5 weeks in growth chamber (Figure 6). Thus, *HvFT3* is expressed in a winter cultivar only after there has been some cold exposure, and increasingly with plant age. It is particularly remarkable that the expression of *HvFT3* was correlated with earlier flowering, although it was detected only after the transition from vegetative to reproductive apex had occurred (Fig. S3a and S4). This late effect on development is consistent with findings in spring wheat varieties (Halliwell *et al.*, 2016), and in a GWAS study in barley (Alqudah *et al.*, 2014). This last study associated polymorphisms at the *HvFT3* region with the time to tipping and the sub-phase awn primordium-tipping.

The induction of *HvFT3* in sowing event 2 (cross-sectional sampling in experiment 1), together with the progressive increase of the transcripts after 28-days VER, when *HvVRN2* is not detected, are consistent with the antagonistic role between *HvVRN2* and *HvFT3* revealed by Casao *et al*., (2011a). However, there were samples in which the absence of *HvVRN2* did not spur the expression of *HvFT3*, showing that the antagonistic relationship is not perfect. These findings overall support that *HvVRN2* absence allows induction of *HvFT3*, but also indicate that it is not sufficient to ensure *HvFT3* expression, hinting at the possible involvement of other repressors. *HvOS2* showed an inverse relationship with *HvVRN1* and *HvFT1* in the cross-sectional sampling (sowings 2, 10 and V in Fig. 4). This finding highlights the interest of further studying the role of *HvOS2*, and its possible relationship with *HvFT3* (already pointed out by Cuesta-Marcos *et al.*, 2015).

*HvFT3* expression was paralleled by that of *HvVRN1* and *HvFT1*. Previous reports agree with our observation. Lv *et al.*, (2014) reported in *Brachypodium* and wheat, that developmental changes regulated by *FT1* were related to transcript levels of other *FT*-like genes, as *FT3*. Under LD, these authors only found upregulation of *FT3* when *FT1* was upregulated, similarly to our findings. In this respect, Li *et al.*, (2015) demonstrated different interactions between FT1 and other FT-like proteins, including FT3, with FD-like and wheat and barley 14-3-3 proteins, all components of the florigen activation complex (FAC). Indeed, they showed a high specificity in the interaction of the protein HvFT3 with Ta14-3-3C, revealing its importance in flowering.

### Coordinated expression of photoperiod and vernalization intermediaries *HvCO2* and *HvVRN2*

*HvVRN2* expression usually occurs in LD. Under SD conditions (8-9 h), the repression of *HvVRN2* is controlled by components of the circadian clock (Turner *et al.*, 2013), although expression under SD, due to the overexpression of *HvCO2*, has been reported (Mulki and von Korff, 2016). Our findings suggest an apparent coordination of *HvCO2* and *HvVRN2* responses in three-week old plants in experiment 1. Up to sowing event 4, the expression of both genes was low, albeit gradually increasing in both cultivars. After that, expression of *HvCO2* dropped dramatically, concurrent with *HvVRN2* raise at sowings 5 and 6. This pattern is consistent with the reported competition between these proteins for binding to NF-Y proteins (Li *et al*. 2011), and also with the feedback loop proposed by Mulki and von Korff (2016). At some point between sowing events 4 and 5, there is a tipping point in expression, possibly related to the dynamics of these two proteins, which could shift the balance of the feedback loop towards higher expression of *HvVRN2*. Then, at event 8 and on, the relationship between the expressions of these two genes seems less tight. Also, at later date, (under 15 h), the relationship presented clear genotypic differences. At that moment, *HvVRN2* expression remained relatively strong (in absence of vernalization); *HvCO2* expression, however, showed a strong recovery after sowing event 4 in ‘Hispanic’, whereas ‘Barberousse’ steadily showed low expression. Therefore, there is a clear shift in the balance of these two genes when day-length is longer than 12h 30min. From that point on, the two genotypes present different patterns.

The control of these two genes has been linked to *Ppd-H1*. Mulki and von Korff (2016)presented evidence of another feedback loop, between *HvVRN2* and *Ppd-H1*, whereas the induction of *HvCO2* by *Ppd-H1*, proposed in the past (as reviewed in Campoli *et al*. 2014), is currently questioned (Chen *et al.*, 2014; Song *et al.*, 2015). In any case, it is clear that both *HvVRN2* and *HvCO2* respond to day-length, either directly through *PHYTOCHROME C* (*PhyC*), or having *Ppd-H1* as an intermediary (as reviewed by Song *et al.*, 2015). *Ppd-H1* (*HvPRR37* as reported by Campoli *et al.*, 2012) is the long photoperiod sensitivity gene in barley, and its expression is under circadian control, with a broad expression peak around 12 h of light in LD (Turner *et al.*, 2005; Campoli *et al.*, 2012). Consequently, its maximum expression levels require days of 12 h or longer. Although sampling times do not match that peak, we can observe and effect over the expression of *HvCO2* and *HvVRN2*, which gradually increased under longer days. The tipping point at 12 h 30 min actually agrees with the date when natural day-length surpasses the maximum expression threshold for *Ppd-H1*. Recently, it was demonstrated in *A. thaliana* that different PRRs not only induce *CO* transcription, but also stabilize the CO protein during the day, enabling to accumulate under LD and initiate floral transition (Hayama *et al.*, 2017). The role of these proteins in cereals should be further clarified.

Mulki and von Korff (2016) proposed that, the dominant *Ppd-H1* could be acting as a flowering repressor before vernalization is fulfilled, which usually takes place under non-inductive photoperiods. We could say that it has a direct impact onto the vernalization requirement of a genotype, as higher *HvVRN1* induction may be needed to down-regulate the increased *HvVRN2* levels brought about by *Ppd-H1* induction, although a parallel mechanism to explain *Ppd-H1* delaying effects in facultative barleys could exist.

### Photoperiod sensitivity through *Ppd-H1* delays field heading/flowering date irrespective of the vernalization process

The mechanism just described sheds light on a phenomenon repeatedly observed in field conditions: flowering delay associated with the dominant (sensitive) allele at *Ppd-H1*, and when field trials contrasted for flowering Julian date. A QTL peak for heading date locating with *Ppd-H1*, with opposite effects in trials, was found in different biparental populations (Ponce-Molina *et al.*, 2012; Mansour *et al.*, 2014). At the earliest trials, the sensitive *Ppd-H1* allele slightly delayed heading, whereas accelerated it in later trials. *HvVRN2* is present in these two populations and, therefore, the delaying effect could be the result of *HvVRN2* expression reinforcement (Mulki and von Korff, 2016). Similar findings were reported for populations Dicktoo × Morex (Pan *et al.*, 1994) and Steptoe × Morex (Borràs-Gelonch *et al.*, 2012), in which *HvVRN2* is absent. A parallel mechanism is needed to explain this effect. In all four populations, the sensitive *Ppd-H1* allele was responsible for delaying heading or flowering dates when it occurred in environments with short day-lengths. It is expected, given the locations used, that in most of these experimental situations there was no lack of natural vernalization. Yet, it is possible that there was a window of opportunity for *Ppd-H1*-dominant genotypes to induce expression of *HvVRN2*, or other repressors, at higher levels than would occur in *Ppd-H1*-recessive genotypes, thus causing its delaying effect.

### Transition to reproductive stage can be achieved through several ways

Traditionally, the transition to the reproductive stage has been associated to expression of *HvFT1*, whose protein is translocated from the leaves to the apices (Song *et al.*, 2015). In our experiments, under lesser-inductive conditions, *HvFT1* expression was not always paralleled by expression of *HvVRN1* (experiment 2). We observed *HvCO2* expression concurrently with induction of *HvFT1*, even in absence of vernalization and, remarkably, without detectable expression of other promoters, such as *HvVRN1* or *HvFT3*. These results agree with the transcriptome data reported by Digel *et al.* (2015), who showed expression of *HvFT1* and *HvCO2*, dependent on *Ppd-H1* in LD, in leaves. These authors found that expression in leaves was higher when the apices had passed the double ridge stage (Waddington stage > 2.0), which could explain the increased transcript levels we found after 49 days under controlled conditions.

Our results suggest that *HvCO2* contributes to *HvFT1* induction, similarly to results reported for wheat (Chen *et al.*, 2014), and is downregulated after vernalization as shown in *B*. *distachyon* (Huan *et al.*, 2013). Li *et al.* (2011) found that both wheat VRN2 and CO2 interacted with the same set of HAP/NF-Y inducer proteins and suggested that both play a role integrating environmental signals for transcriptional regulation of *FT1*. In Arabidopsis, *CO* induces *FT* in LD, and consequently flowering, probably through interaction with its promoter (Andrés and Coupland, 2012). In the winter barley varieties employed in this study, under non-inductive LD conditions (Experiment 1, points 6-10, Fig. 2 and Fig. 4), the expression profile of both *CONSTANS* genes (*HvCO2*, *HvCO9*), was rather similar, suggesting a common regulation, possibly in an age or photoperiod dependent-manner.

*HvCO9* is another CCT domain gene, like *HvCO2* or *HvVRN2* (Higgins *et al.*, 2010; Cockram *et al.*, 2012; Kikuchi *et al.*, 2012). Kikuchi et al. (2012) found higher expression of *HvCO9* under SD (12 h light), in spring barleys lacking *HvVRN2*. Our study, using winter barley genotypes, also suggest a repressor role of this gene, stronger in ‘Barberousse’ than in ‘Hispanic’ under SD conditions, related to development and amount of vernalization.

### *HvOS2* could explain differences in development among winter varieties

In this study, striking differences between genotypes have been observed under non-inductive conditions (no or reduced vernalization and SD). ‘Hispanic’ flowered always earlier than ‘Barberousse’, as also evidenced by the expression of all flowering promoters in the first and high expression of the repressors in the latter. The results illustrate the different behavior of these winter genotypes.

The third repressor of flowering time studied, *HvOS2*, showed slight differences between NV and VER plants in early development. The difference became more evident in older plants. Expression of *HvOS2* was highly correlated with the absence of *HvVRN1*, being low in plants which flowered, as was observed in barley, wheat and *Brachypodium* (Greenup *et al.*, 2010; Sharma *et al.*, 2017). Deng *et al*., (2015) showed that the protein VRN1 binds to the promoters of *VERNALIZATION2* and *ODDSOC2*. By now, there is enough evidence substantiating that expression of *OS2* genes in winter cereals is suppressed by cold. For *Brachypodium*, it has been proposed that *BdODDSOC2* “plays a role in setting the length of the vernalization requirement in a rheostatic manner, i.e. higher *ODDSOC2* transcript levels before cold result in a longer cold period needed to saturate the vernalization requirement” (Sharma *et al*., 2017), although its specific role in the vernalization response is not clear.

In our results, variety ‘Barberousse’ showed higher levels of *HvOS2* transcripts than ‘Hispanic’ at most sampling times. This higher expression in ‘Barberousse’ is consistent with its delay in development compared to ‘Hispanic’.

### *HvVRN1* expression in winter barley under non-inductive conditions occurred after the apex transition

‘Hispanic’ and ‘Barberousse’ showed differences in development in response to photoperiod and vernalization. The overall expression patterns for flowering genes were in line with expectations for winter varieties. Commonly, winter varieties need to undergo a specific number of cold hours before flowering. The varieties used in this study carry *vrn1* allele, with a full-length intron and present a strong vernalization need (Fig. S1). In absence of vernalization, there is no expression of *HvVRN1* and consequently, flowering is delayed, as expected (Trevaskis *et al.*, 2003; Yan *et al.*, 2003; von Zitzewitz *et al.*, 2005). We did not observe expression of *HvVRN1* in NV 21-day-old plants at any point. Only plants from sowing event 2, 13 weeks after sowing, showed *HvVRN1* expression. This result agrees with previous reports of the induction of *HvVRN1* by development after 10-12 weeks in unvernalized winter plants, under LD (Trevaskis *et al.*, 2006). The lack of *HvVRN1* expression in non-vernalized plants is related to the presence of *HvVRN2* and *HvOS2* transcripts, as was evidenced by other authors (Dubcovsky *et al.*, 2006; Trevaskis *et al.*, 2006; Greenup *et al.*, 2010; Deng *et al.*, 2015; Sharma *et al.*, 2017). When transcripts of these flowering repressors were present, the flowering promoters *HvVRN1*, *HvFT1* and *HvFT3* were directly or indirectly downregulated, and flowering was delayed (Fig. 4, Fig. 6). Indeed, Deng *et al.* (2015) showed that, in winter, VRN1 controls directly *HvFT1* levels binding to its promoter, and indirectly through *HvVRN2* and *HvOS2* down-regulation. This repression could also be stimulating *HvFT3* and expression of other flowering promoters (Cuesta-Marcos *et al.*, 2015).

We observed a more loose relationship between *HvVRN1* and *HvFT1* than expected. Some apices developed to reproductive stage (W2, Fig. S4), even though this was not paralleled by induction of *HvVRN1* in the leaves at the same time in both experiments (Fig.4, Fig.6). Indeed, the peak of expression of *HvVRN1* seems to be related with the appearance of the floret primordium (W3.5). While this result could be in conflict with the essential role of *HvVRN1* in the initiation of reproductive phase described before (Trevaskis *et al.*, 2003), data by Digel *et al.* (2015) can clarify our observations. These authors showed that *HvVRN1*, together with other MADS box transcription factors, was upregulated in the leaves and shoot apices during pre-anthesis, but the transcript levels of *HvVRN1* were first induced in the shoot apices. This gap would explain why we did not detect *HvVRN1* expression in the leaves although some apices had already progressed in their development.

### Conclusion

The use of different sowing events, under natural increasing photoperiod corroborate that the expression of *HvVRN2* is highly dependent on day-length, and we provide evidence of the threshold, around 12 h 30 min, above which this expression rises markedly and affects most plant development. This experiment also highlighted the importance of completing the vernalization requirement before a certain day-length threshold, in order to promote flowering in optimum conditions. *HvFT3*, a central gene for winter barley performance in Southern Europe, is not induced just by short days. In winter cultivars with dominant *Ppd-H1*, it must receive additional induction through either the autonomous pathway, and/or a cold period, to be effective in reducing time to flowering.

In winter barleys, *HvVRN2* transcript levels are always present, but we propose that its activity (and that of *HvOS2*) must be below a functional threshold to allow timely flowering, which will not occur in absence of vernalization. Here, we emphasize the importance of *HvVRN2* in the promotion to flowering, but also the role of *HvOS2* and *HvCO2* in vernalization-responsive cultivars. *HvOS2* seems to contribute to *HvVRN2* function in the delay of flowering, while *HvCO2* might be promoter of *HvFT1* in both inductive and non-inductive conditions, being affected by those two repressors and *HvCO9*.

The photoperiod conditions of the experiments here described, correspond to a wide range of late spring sowings for winter barley in the Mediterranean area. The genetic mechanisms and the environmental controls involved in this study will be useful to define both varieties and agronomics best suited for current and future climate conditions.

## Supplementary data

Supplementary Table 1. Primer sequences.

Supplementary Figure S1. Flowering date under different vernalization treatments in the preliminary experiment.

Supplementary Figure S2. Gene expression in 2-week-old plants sown under natural and increasing photoperiods (without vernalization and control).

Supplementary Figure S3. Associations between gene expressions and heading date (DEV49) in experiment 2.

Supplementary Figure S4. Apex development in experiment 2 (growth chamber, 12 h day).

## Acknowledgements

Study financially supported by the Spanish Ministry of Economy, Industry and Competitiveness (Projects AGL2013-48756-R, including a scholarship granted to AM, and AGL2016-80967-R).

